# ePlatypus: an ecosystem for computational analysis of immunogenomics data

**DOI:** 10.1101/2022.09.28.509709

**Authors:** Victor Kreiner, Andreas Agrafiotis, Tudor-Stefan Cotet, Raphael Kuhn, Danielle Shlesinger, Marcos Manero-Carranza, Keywan Khodaverdi, Solène Massery, Lorenzo Guerci, Kai-Lin Hong, Jiami Han, Kostas Stiklioraitis, Vittoria Martinolli D’Arcy, Raphael Dizerens, Samuel Kilchenmann, Lucas Stalder, Leon Nissen, Basil Vogelsanger, Stine Anzböck, Daria Laslo, Melinda Kondorosy, Marco Venerito, Alejandro Sanz García, Isabelle Feller, Annette Oxenius, Sai T. Reddy, Alexander Yermanos

**Affiliations:** Department of Biosystems Science and Engineering, ETH Zurich, Basel, Switzerland; Institute of Microbiology, ETH Zurich, Zurich, Switzerland; Department of Pathology and Immunology, University of Geneva, Geneva, Switzerland; Center for Translational Immunology, University Medical Center Utrecht, Utrecht, Netherlands

## Abstract

The maturation of systems immunology methodologies requires novel and transparent computational frameworks capable of integrating diverse data modalities in a reproducible manner. Here, we present the ePlatypus computational immunology ecosystem for immunogenomics data analysis, with a focus on adaptive immune repertoires and single-cell sequencing. ePlatypus is a web-based platform and provides programming tutorials and an integrative database that elucidates selection patterns of adaptive immunity. Furthermore, the ecosystem links novel and established bioinformatics pipelines relevant for single-cell immune repertoires and other aspects of computational immunology such as predicting ligand-receptor interactions, structural modeling, simulations, machine learning, graph theory, pseudotime, spatial transcriptomics and phylogenetics. The ePlatypus ecosystem helps extract deeper insight in computational immunology and immunogenomics and promote open science.

**Accessibility:** https://alexyermanos.github.io/Platypus/index.html

## Main

The fields of systems and computational immunology have advanced substantially in recent years, most notably through progress in genomics and single-cell sequencing, which are transforming the measurement of adaptive immune responses from qualitative to quantitative science. In recent years, a number of bioinformatic software tools have been developed that provide rapid and facile exploration of single-cell RNA sequencing (scSeq) data and perform analysis such as differential gene expression, cell clustering and transcriptional phenotyping (Efremova et al. 2020; Satija et al. 2015). However, in the context of immunogenomics, lymphocytes (B and T cells) and their transcriptomes and immune receptor repertoires (B cell receptor, BCR and T cell receptor, TCR), there is a lack of software enabling the simultaneous interrogation and integration of multiple approaches capable of deconstructing high-dimensional immune responses, such as phylogenetics, machine learning, graph theory, and structural modeling. Moreover, although deep sequencing of immune repertoires has become a common method in modern immunology, locating, downloading, and integrating data across experiments and research groups remains challenging. Finally, most immunogenomics software tools require computational expertise involved in analyzing such feature-rich datasets (Borcherding et al., 2020; Yaari & Kleinstein, 2015; Yermanos et al., 2021).

Here, we present ePlatypus, a computational immunology ecosystem that expands upon Platypus, a previously developed immunogenomics software. The ePlatypus ecosystem (Figure 1) consists of an R package hosted on CRAN and possesses hundreds of functions, including those most relevant for single-cell immunogenomics (transcriptome and immune repertoires) as well as many other aspects of computational immunology. These include the following: i) profiling the relationship between clonal expansion and transcriptional phenotypes to identify distinct activation phenotypes of expanded lymphocytes (Figure S1), ii) pseudobulk differential expression pipelines to robustly characterize transcriptional clusters leveraging methods originally designed for bulk RNA-sequencing (Figure S2), iii) immune repertoire diversity metrics to characterize clonal distributions and to ensure sufficient sampling depths have been recovered (Figure S3), iv) phylogenetics to identify evolutionary trajectories and intraclonal network properties of B cells during infection (Figure S4), v) B and T cell sequence similarity networks to identify fundamental principles of lymphocyte repertoire architecture in the course of an immune response (Figure S4), vi) machine-learning guided classification to predict BCR and TCR specificity and further uncover feature importance of antigen-specific sequences (Figure S5), vii) predicting ligand-receptor interactions under homeostatic and disease conditions using the CellphoneDB repository (Efremova et al. 2020) (Figure S6), viii) spatial transcriptomics to spatially interrogate gene expression patterns and further integrate clonal selection and clonal evolution of adaptive immune responses (Figure S7), ix) and structural modeling of immune receptor sequences and repertoires using multiple external tools including AlphaFold, IgFold, and DeepAb (Ruffolo et al. 2022; Jumper et al. 2021) (Figure S8). Importantly, ePlatypus currently hosts an online portal with 20 tutorials and walk-throughs (Figure S9), each of which contain code, comments, and explanatory text (Figure S10) for various computational immunology frameworks (Table S1, Figure S1).

**Figure 1.**
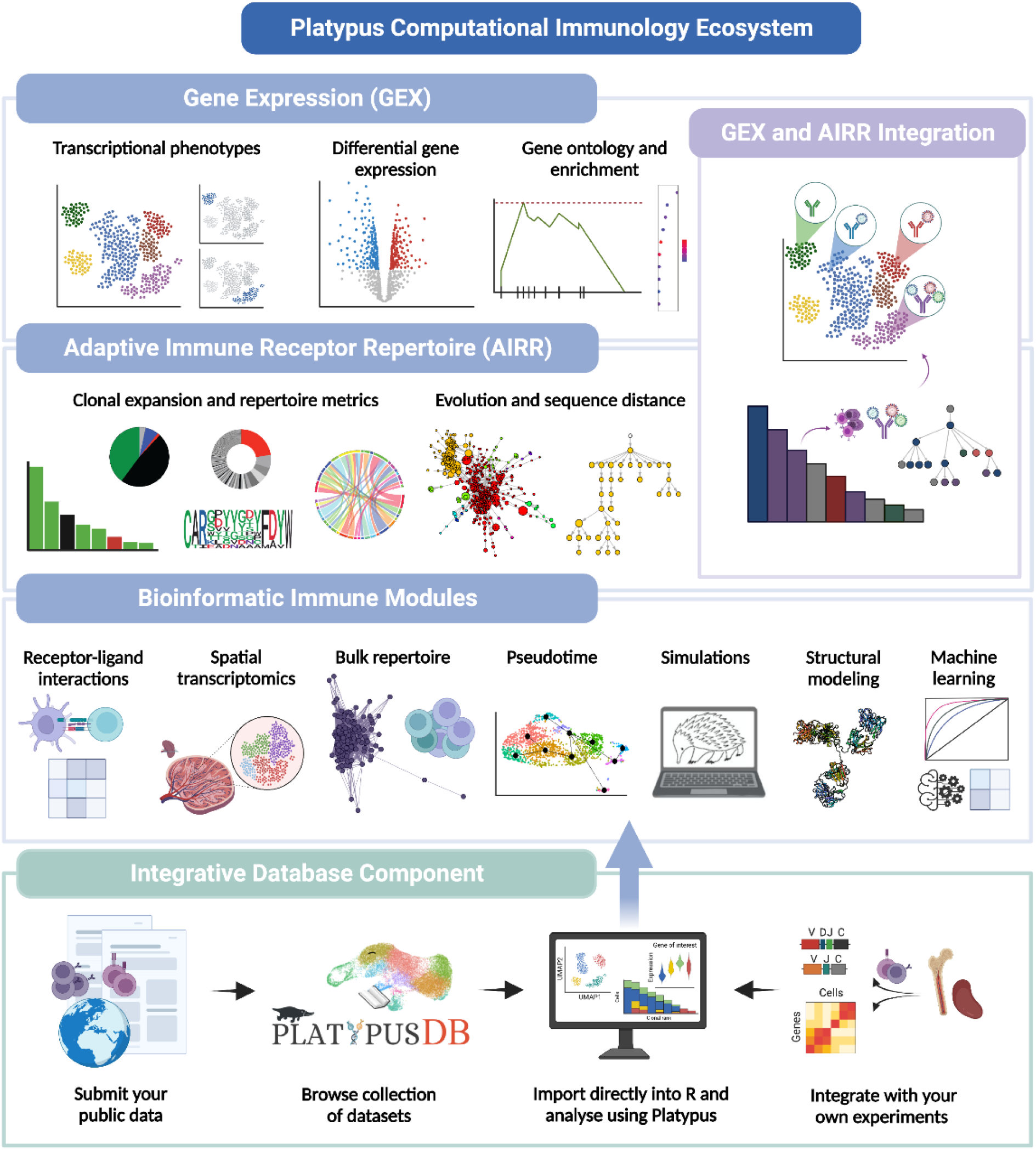
Breadth of the ePlatypus Computational Immunology Ecosystem. The ecosystem currently is composed of a core R package that has pipelines pertaining to immune repertoires, gene expression, receptor-ligand interactions, spatial transcriptomics, pseudotime, simulations, structural modeling and machine learning. Similarly, the ecosystem contains an integrated database and a website currently containing 20 tutorials with accompanying code.

Additionally, the ePlatypus ecosystem contains a database component, PlatypusDB, that directly integrates into the R programming language, thereby allowing the rapid analysis and integration of B and T cells containing both adaptive immune receptor information (VDJ) and single-cell transcriptomes (GEX). PlatypusDB both stores raw output files from the commonly used aligner tool Cellranger (10x Genomics) and also holds the immune-relevant data in the form of an R object that can be loaded directly into the R environment without explicitly requiring file download. Importantly, the data is stored as both the processed aligned output and as a preprocessed R object that contains transcriptome, immune repertoire, and metadata information. Within the programming interface, the user has the ability to perform the following actions: i) download entire public sequencing datasets, ii) download individual samples from publications, and iii) download and integrate public repertoires with samples stored locally (Figure 1). While the ePlatypus development team will continuously update the ecosystem with newly published datasets, external users can also submit their preprocessed immune receptor repertoires directly for manual curation and addition to the database.

To demonstrate several use cases of the ePlatypus computational ecosystem, we integrated and analyzed multiple single-cell transcriptomes and immune receptor repertoires across different disease conditions, viral infections, and vaccination studies. We directly downloaded murine T cell repertoires from previously published datasets containing both CD4 and CD8 T cells from conditions such as acute and chronic viral infections (Khatun et al., 2021; Kuhn et al., 2022; Merkenschlager et al., 2021; Shlesinger et al., 2022), homeostatic aging (Yermanos, et al., 2021) and experimental autoimmune encephalomyelitis (Shlesinger et al., 2022) (Table S2). Following transcriptional integration with Harmony (Korsunsky et al., 2019), which aims to reduce batch effects across different datasets, we visualized all cells using uniform manifold approximation projection (UMAP) (Figure S11A, S11B). This demonstrated two major transcriptional regions, dominated by either *Cd4* or *Cd8* gene expression, which could be simultaneously interrogated with other known gene markers of activation or exhaustion such as *Cd44, Ifng, Pdcd1*, *Lag3* and *Il7r* (Figures S11B, S12). Supplementing this focused analysis with ProjectTILS, a recently developed reference atlas which helps resolve murine T cell heterogeneity of tumor-infiltrating T cells (Andreatta et al., 2021), demonstrated that T cells from PlatypusDB almost entirely cover the ProjecTILs main reference dataset (Figures S11C, S11D, S11E, S13). To highlight the potential to link repertoire features with transcriptional heterogeneity, we visualized the most expanded T cell clones on the transcriptional landscape (Figure S11F). This demonstrated diverse levels of clonal expansion within the database, with those expanded clones corresponding to a relative upregulation of activation markers compared to those with lower expansion (Figure S12).

Next, we used ePlatypus to investigate whether similar transcriptional heterogeneity could be detected for B cells present in the PlatypusDB. Multiple datasets derived from murine models of infection, immunization and autoimmune disease (Agrafiotis et al., 2021; Mathew et al., 2021b; Neumeier et al., 2021; Shlesinger et al., 2022; Yewdell et al., 2021) were integrated as previously described (Table S2). Transcriptional clustering suggested that common B cell clusters were present across multiple datasets (Figure S14A), which exhibited varying expression levels of markers relating to antibody secretion and B cell differentiation (e.g. *Cd138, Xbp1*, *Slamf7*) (Figure S15) and diverse isotype usage (Figure S14B). The analyses presented here highlights the breadth of B and T cell phenotypes and selection patterns already available within ePlatypus, which will only continue to grow as more user-supplied public datasets are added.

The maturation of systems immunology methodologies requires novel and transparent computational frameworks capable of integrating diverse data modalities in a reproducible manner. The ePlatypus ecosystem, composed of a core R package with hundreds of functions, programming tutorials, and a comprehensive database, helps extract deeper insight in immunogenomics while promoting open science.

---

**Figure S1.**
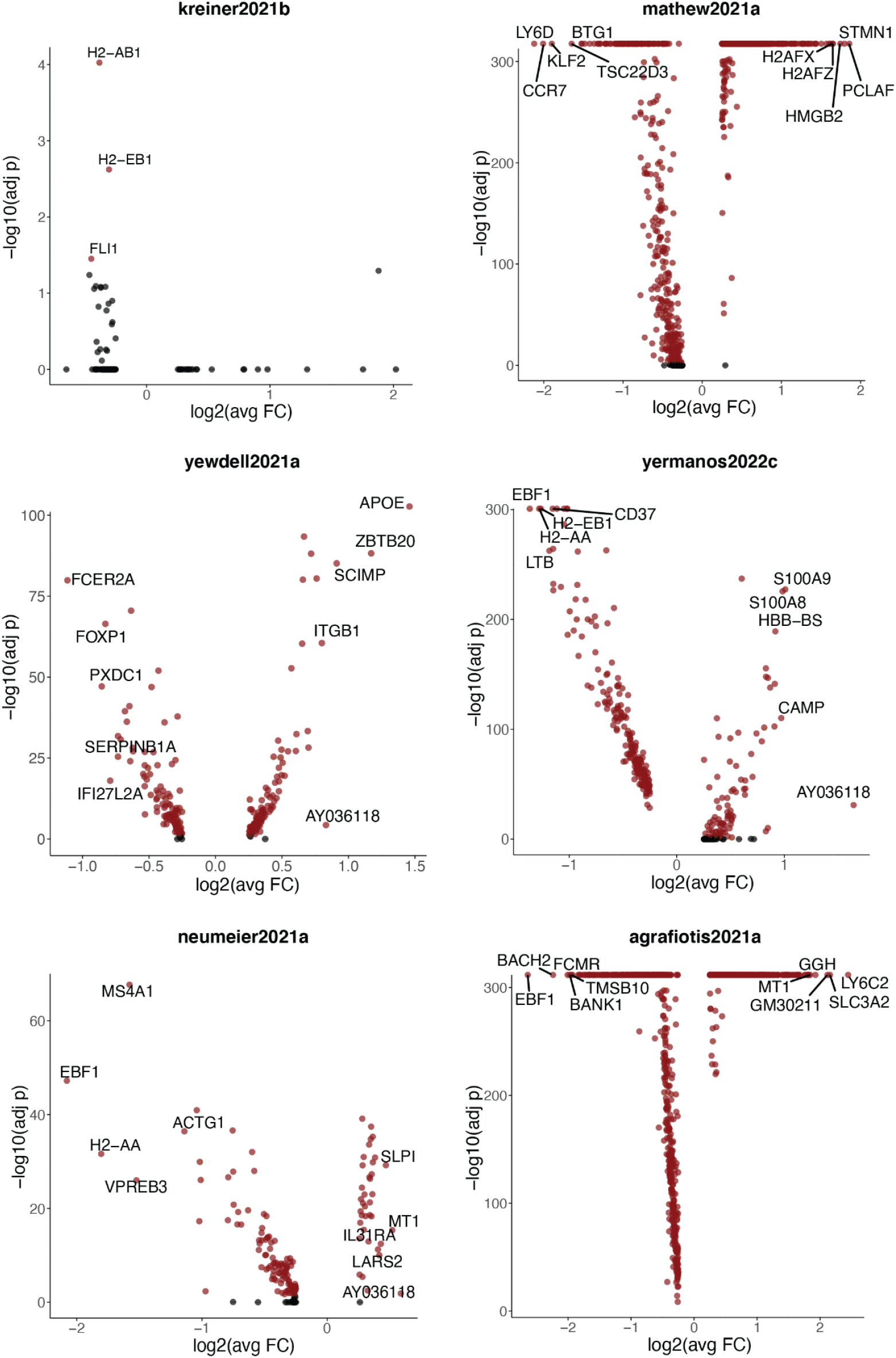
Differentially expressed genes between lowly (n=1 cell per clone) and highly (n>1 cell per clone) expanded clones for different B cell datasets. Positive log2(avgFC) indicates upregulated genes in the expanded clones for all datasets. Headers indicate the origin of the dataset as can be found in PlatypusDB.

**Figure S2.**
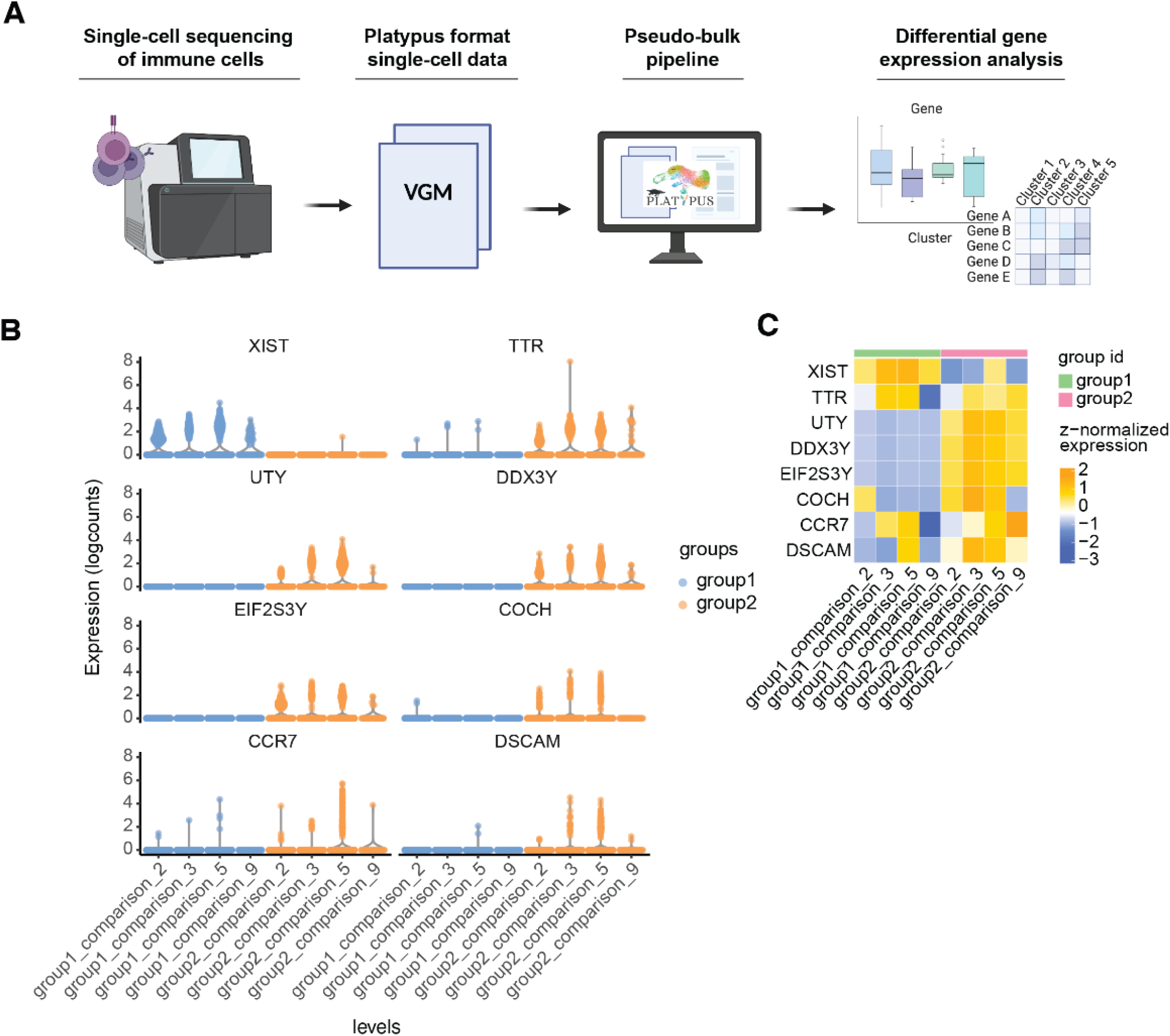
Pseudobulk pipeline within the ePlatypus ecosystem. A. Graphical overview of the Pseudobulk workflow. B. Violin plots displaying the expression levels of differentially expressed genes between two user-defined categories (group 1 and group 2). Single-cell transcriptional clusters 2, 3, 5 and 9 were grouped in comparison levels (or clusters) according to user-defined categories. Cells within the same comparison level belonging to different groups were tested for differential gene expression using the Platypus pseudo_bulk_DE function. C. Heatmap highlighting normalized expression levels of differentially expressed genes between the different comparison levels.

**Figure S3.**
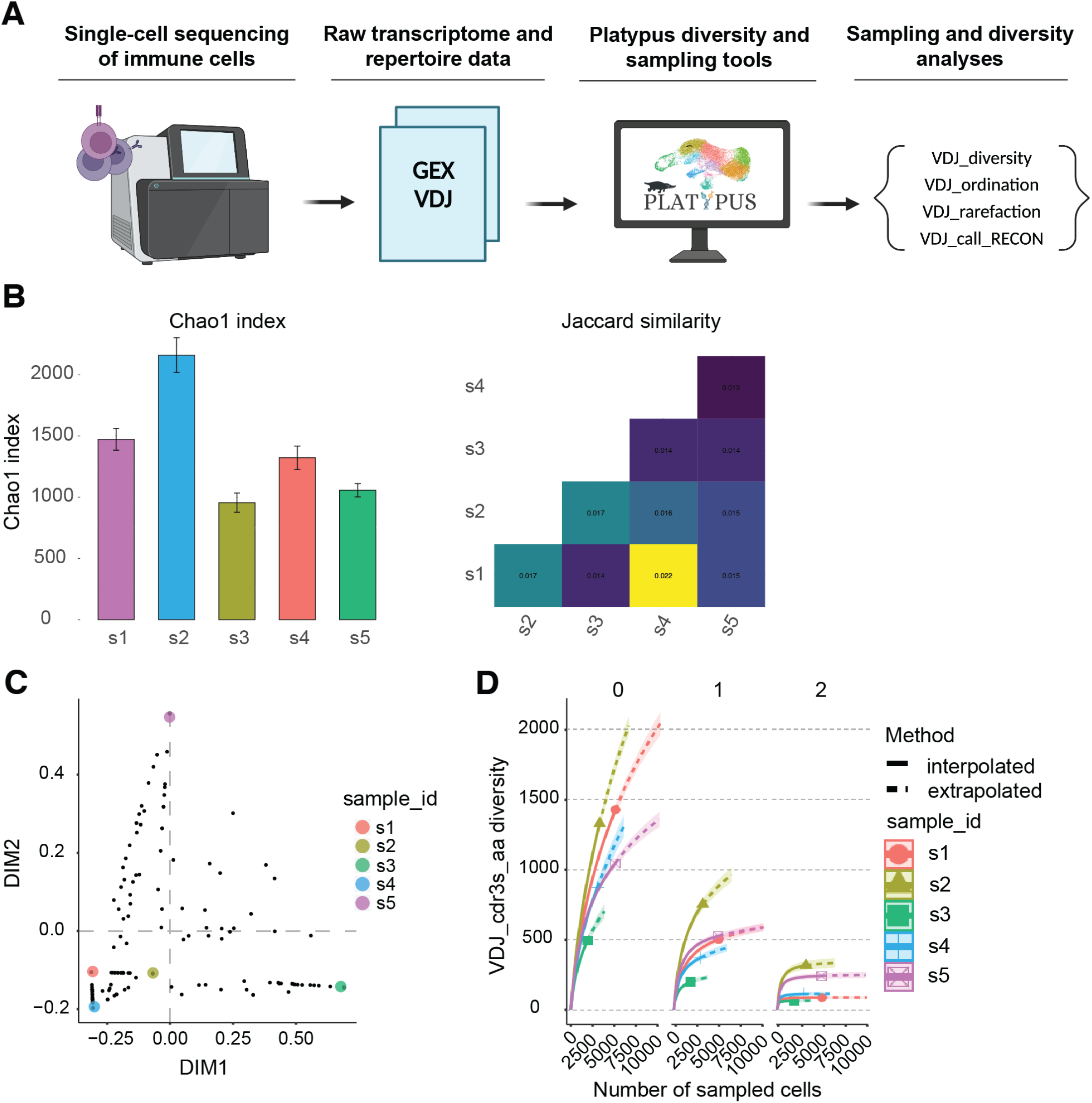
Sampling and diversity analysis pipelines. A. Graphical overview of the sampling and diversity workflow. Single-cell immune repertoire sequencing samples are formatted into a single VDJ_GEX_matrix object, which can be supplied to downstream calculations of diversity, rarefaction, and ordination analyses. B. VDJ_diversity includes a wide range of α and β-diversity calculators. α-diversity is displayed as per-sample bar plots for the Chao1 index, whereas β-diversity is shown as a heatmap. C. Dimensionality reduction on the abundance data can be performed using VDJ_ordination – several dimensionality reduction algorithms are available inside the function (UMAP, t-SNE, PCA, PCoA). D. Rarefaction curves can be displayed using the VDJ_rarefaction function: in this example, the curve indicative of each Hill number is grouped by sample.

**Figure S4.**
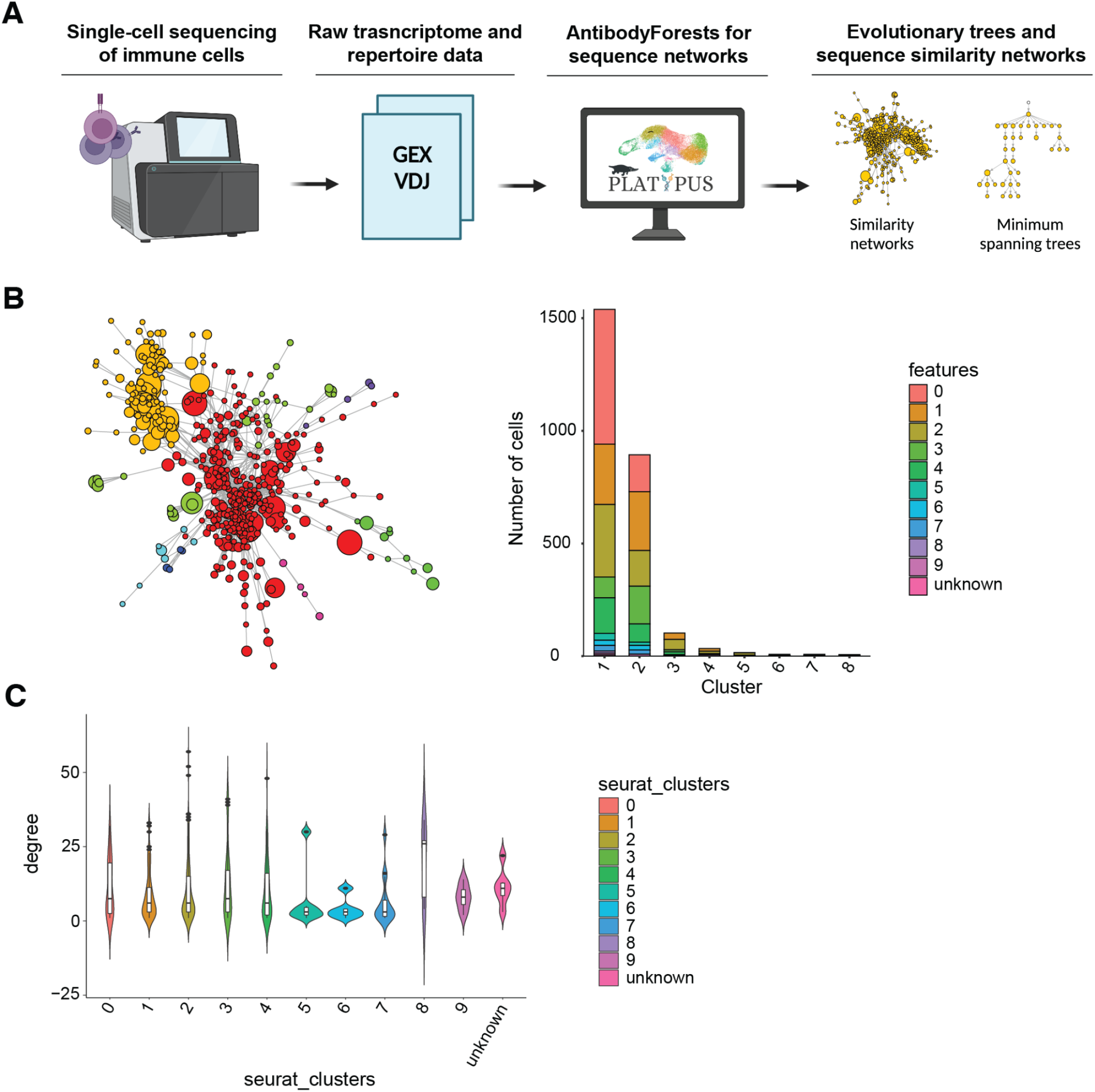
Repertoire similarity networks and quantification. A. The AntibodyForests pipeline designed for network inference, direct visualization, and network analysis of immune receptor minimum spanning trees or sequence similarity graphs. B. Clusters can be determined in a sequence similarity network using the AntibodyForests_communities function and then colored using the AntibodyForests_plot function. Moreover, the tool can produce per-cluster bar plots of various sequence or cell features (e.g., the Seurat transcriptomic cluster per sequence-similarity clusters). C. The AntibodyForests_metrics function calculates a wide range of node and edge metrics for the resulting AntibodyForests networks: in this case, a violin plot depicts the node degree distribution across each Seurat transcriptomic cluster for the similarity graph showcased in B (each node was assigned the most frequent Seurat cluster across all cells with that particular receptor sequence).

**Figure S5.**
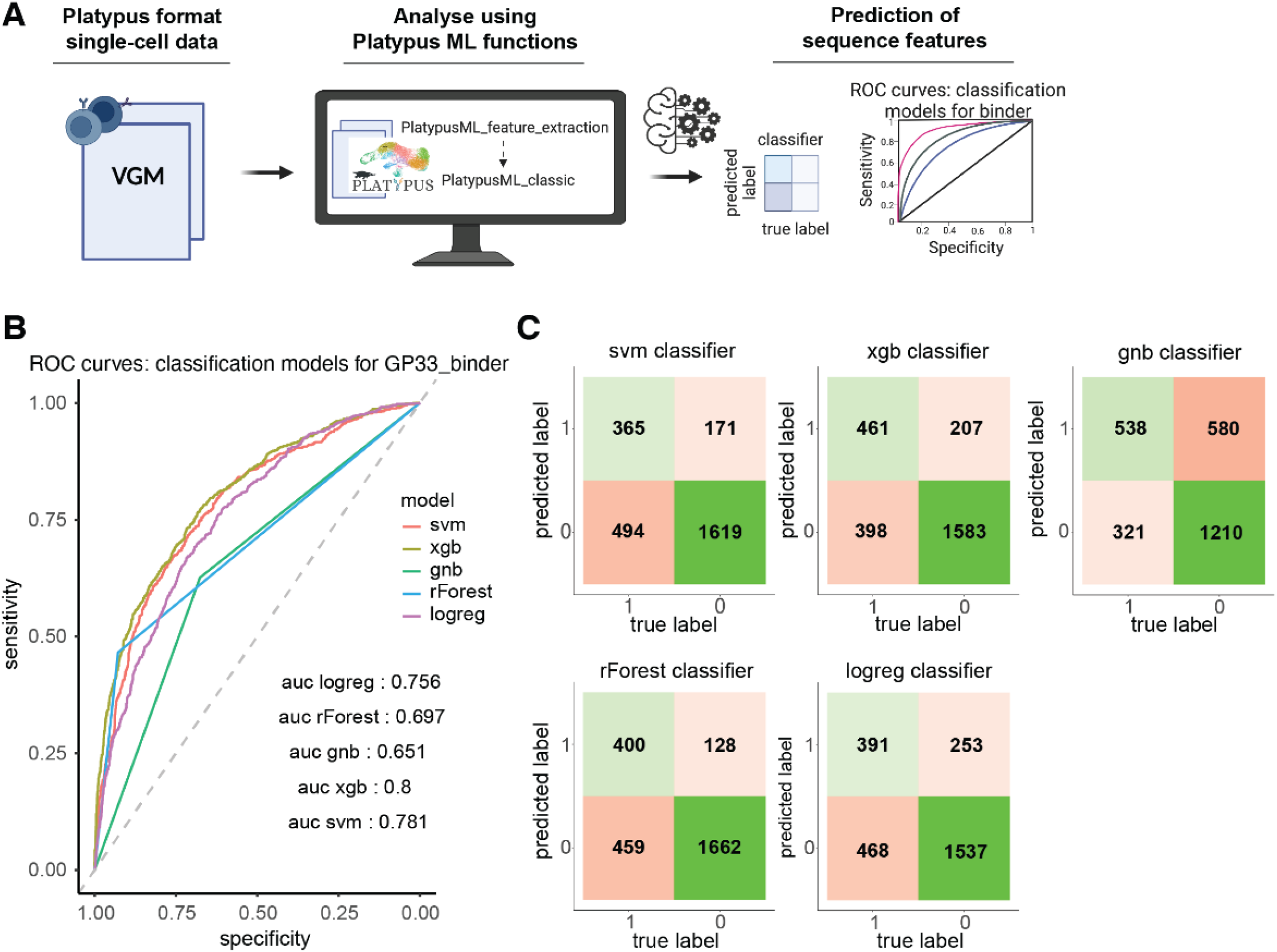
Machine learning pipelines for adaptive immune receptor classification. A. Workflow for machine learning (ML) classification of virus-specificity within the ePlatypus Ecosystem. B. Receiver operating characteristic (ROC) curves and area under the curve (AUC) scores for different classification models (logreg: logistic regression, rForest: random forest, gnb: gaussian naive bayes, xgb: XG Boost, svm: Support vector machine) for binary binding vs non-binding classification of virus-specific T cell receptors following viral infection. C. Confusion matrices for each of the models shown in (B). Green intensity is proportional to the number of true positives and true negatives, while red intensity is proportional to the number of false negatives and false positives.

**Figure S6.**
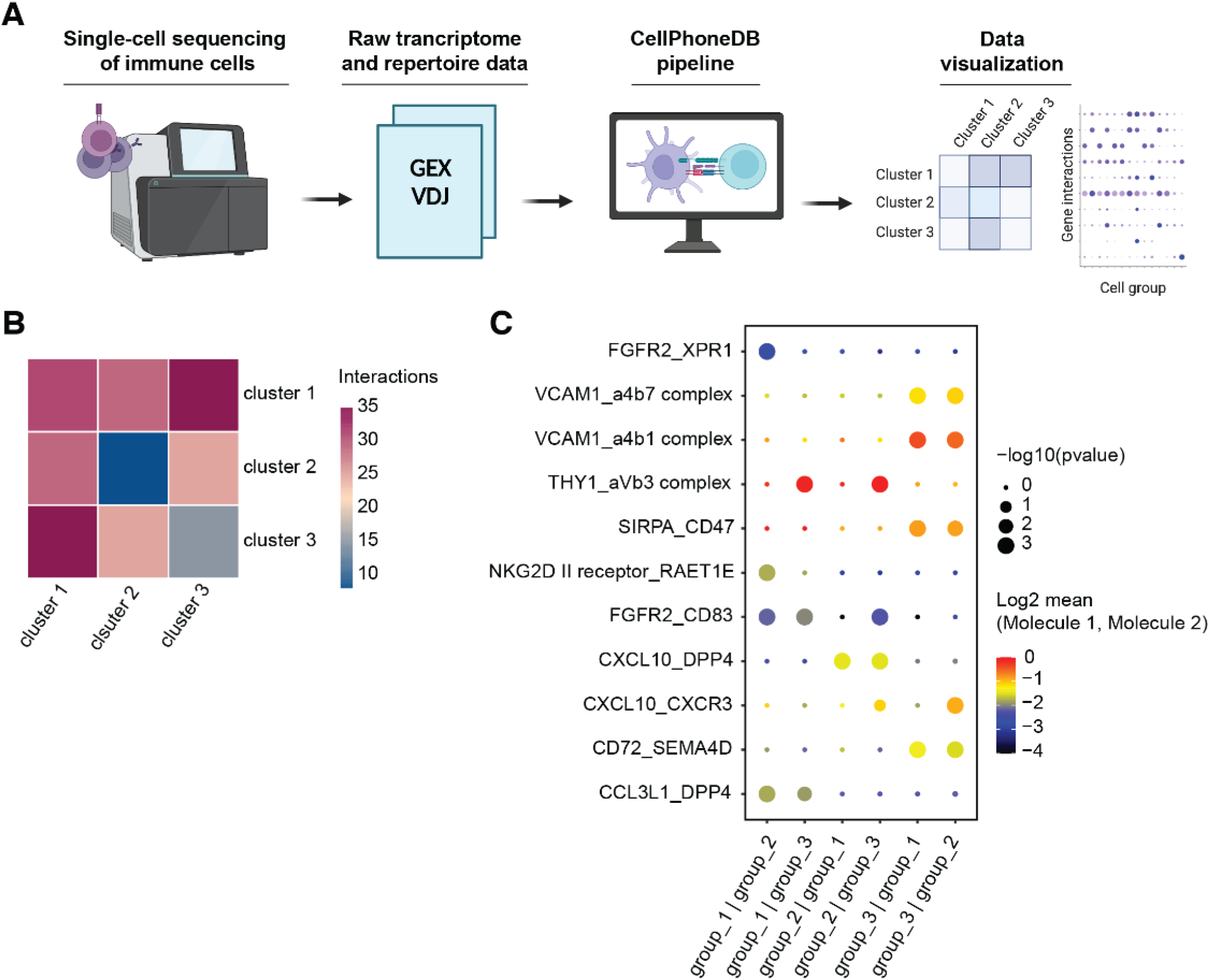
Predicting receptor-ligand interaction using CellphoneDB. A. Graphical overview for processing, generating, and analyzing ligand-receptor interactions. B. Heatmap displaying the level of interaction between cells belonging to different clusters. The interaction level is calculated on the single-cell gene expression data using the CellPhoneDB software. The heatmap is part of the output from the Platypus CellPhoneDB_analyse function. C. Dot plot depicting the interaction levels of the gene pairs reported (y-axis) in the cluster pairs reported (x-axis). Color of the dot indicates the logarithm of the ligand-receptor interaction mean value across cells of the cluster pair. Size of the dot indicates the negative logarithm of the p-value, highlighting which interactions are to be considered significant. The dot plot is part of the output of CellPhoneDB_analyse. A customized version of the plot can be generated using the dot_plot function by selecting the gene and cluster pairs to be displayed, in addition to a mean or p-value threshold.

**Figure S7.**
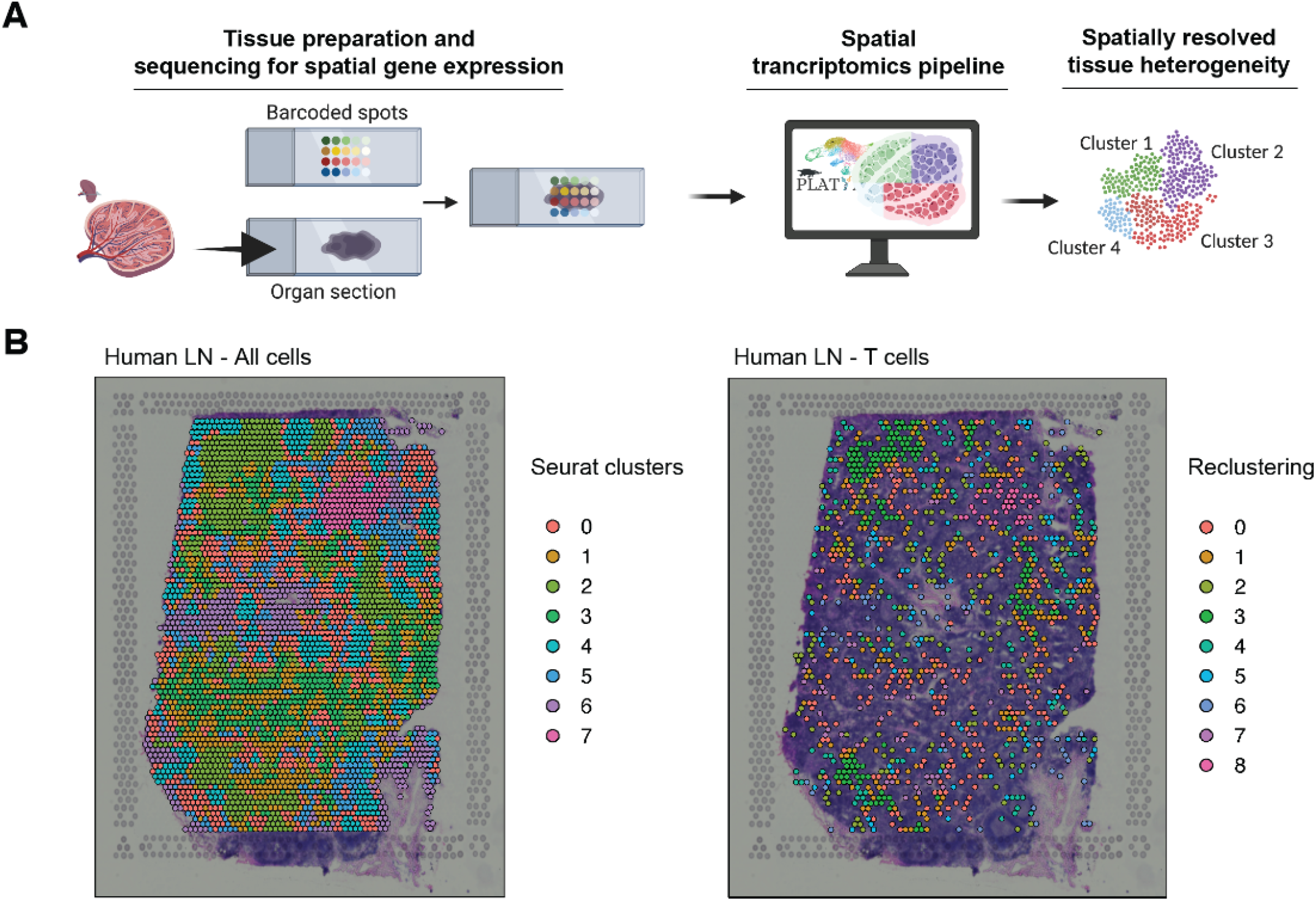
Spatial transcriptomics analysis pipeline. A. Graphical overview for the generation and analysis of spatial transcriptomics experiments. B. Unsupervised transcriptional clustering of cells from a human lymph node (LN) sample. The colors indicate the cluster to which the cell belongs for either all cells (right) or only T cells (right).

**Figure S8.**
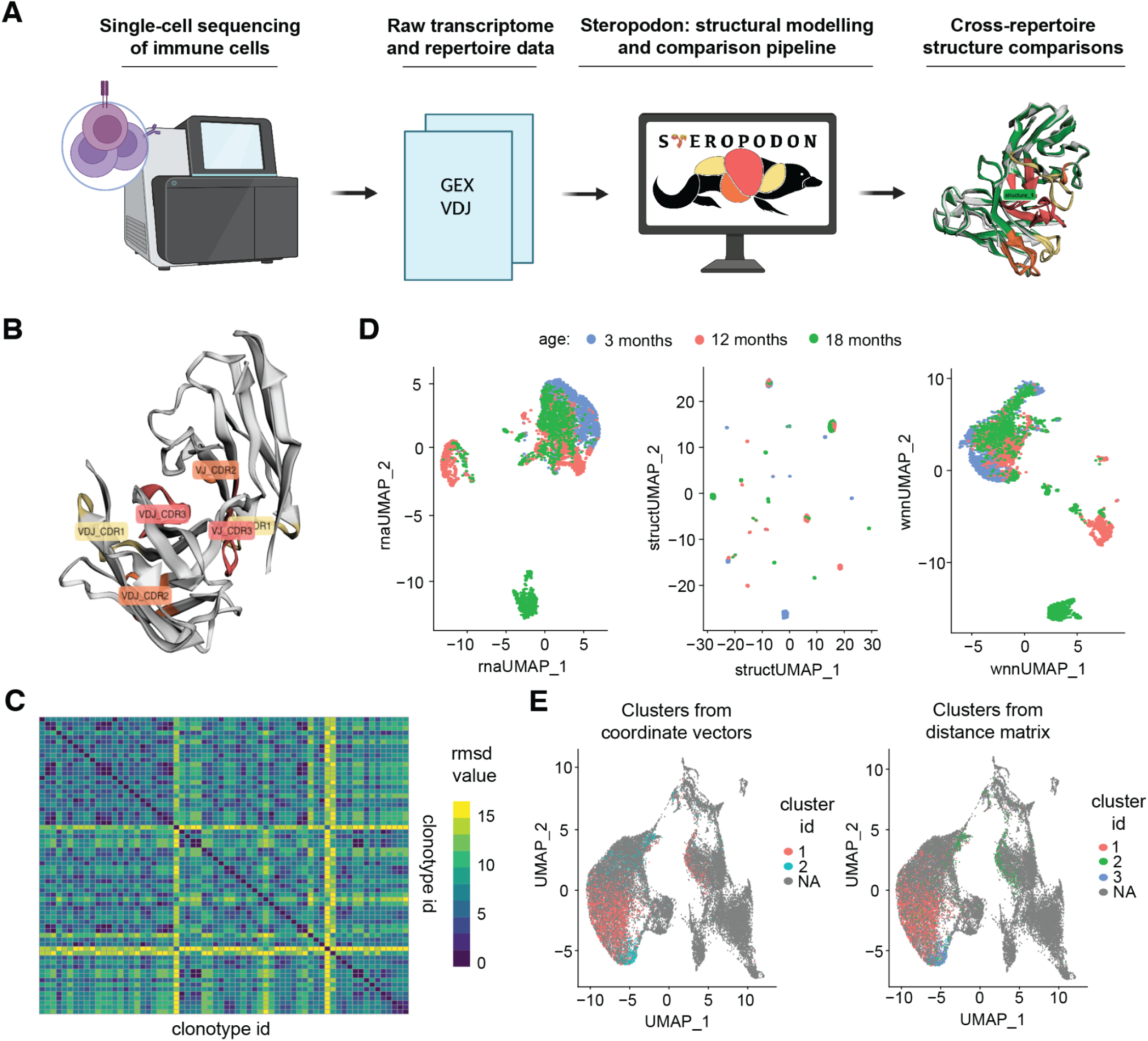
The Steropodon pipeline for immune receptor structural modeling and integrated, cross-repertoire analysis. A. The Steropodon workflow for obtaining receptor structures, starting from the Platypus VGM object. B. A sample structure from the TNFR2 dataset obtained via Steropodon_model and visualized using Steropodon_visualize, with the CDR regions labelled. C. A heatmap of the root mean square deviation (RMSD) between structures from the 10 most frequent clonotypes in 6 samples from the TNFR2 dataset, showcasing the overall structural similarities at the intra and cross-repertoire levels. D. UMAP projection of the gene expression vectors for each cell corresponding to the structures modeled from the 10 most expanded clonotypes from the TNFR2 dataset (left), UMAP projection of the coordinate vectors of the modeled structures after structural alignment (middle), and UMAP projection of the multimodal embeddings for the 2 modalities from before, integrated using Seurat’s weighted nearest neighbors algorithm. E. UMAP projection of the GEX values of all cells in the 6 samples selected from the TNFR2 dataset, with cells highlighted by the structural clusters obtained on the coordinate feature vectors from *D* and the distance matrix from *C*, respectively, following hierarchical clustering.

**Figure S9.**
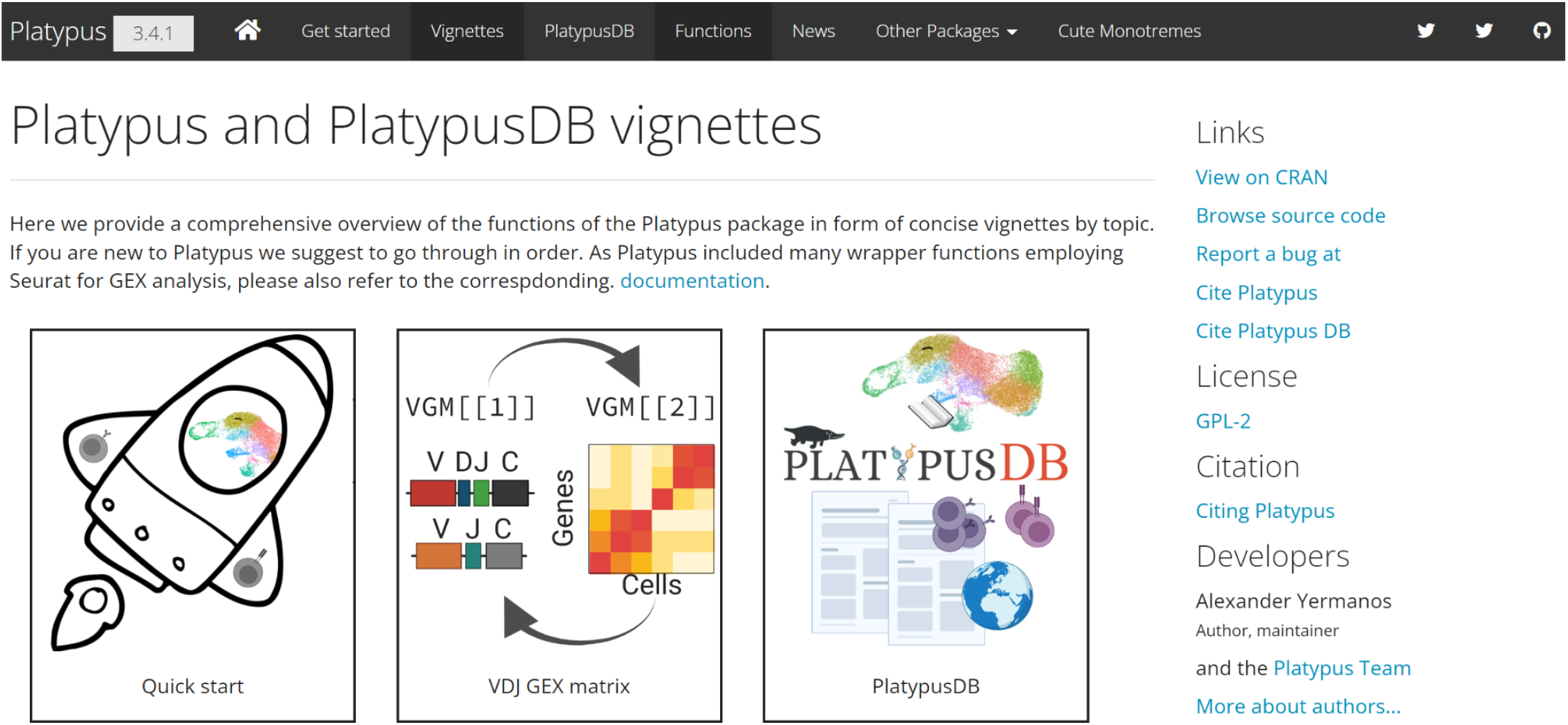
Example of three vignettes available within the ePlatypus Computational Immunology Ecosystem. All other available vignettes presented on the website are detailed in Table S1.

**Figure S10.**
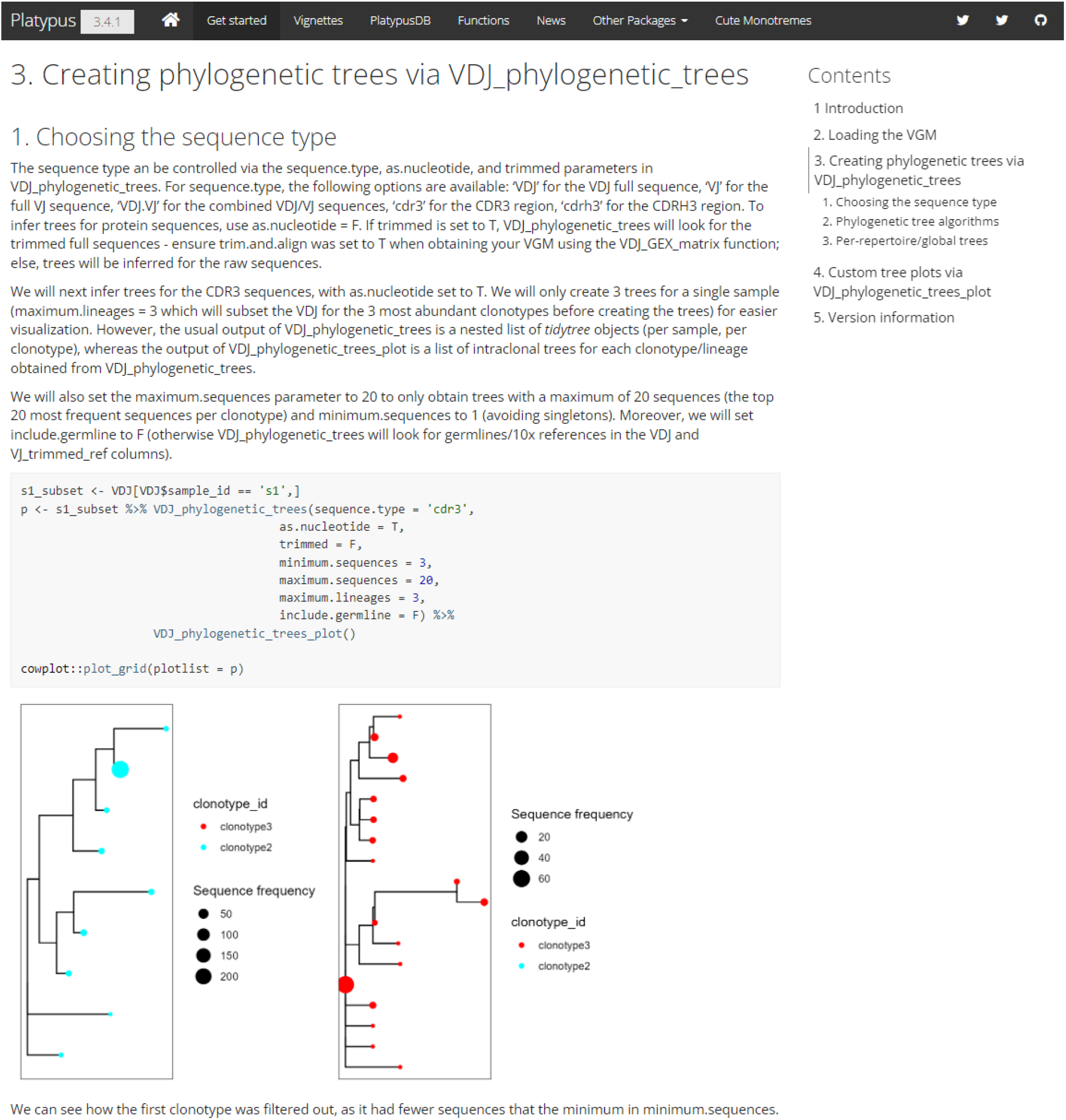
Example of one walk-through present in the ePlatypus Computational Immunology Ecosystem.

**Figure S11.**
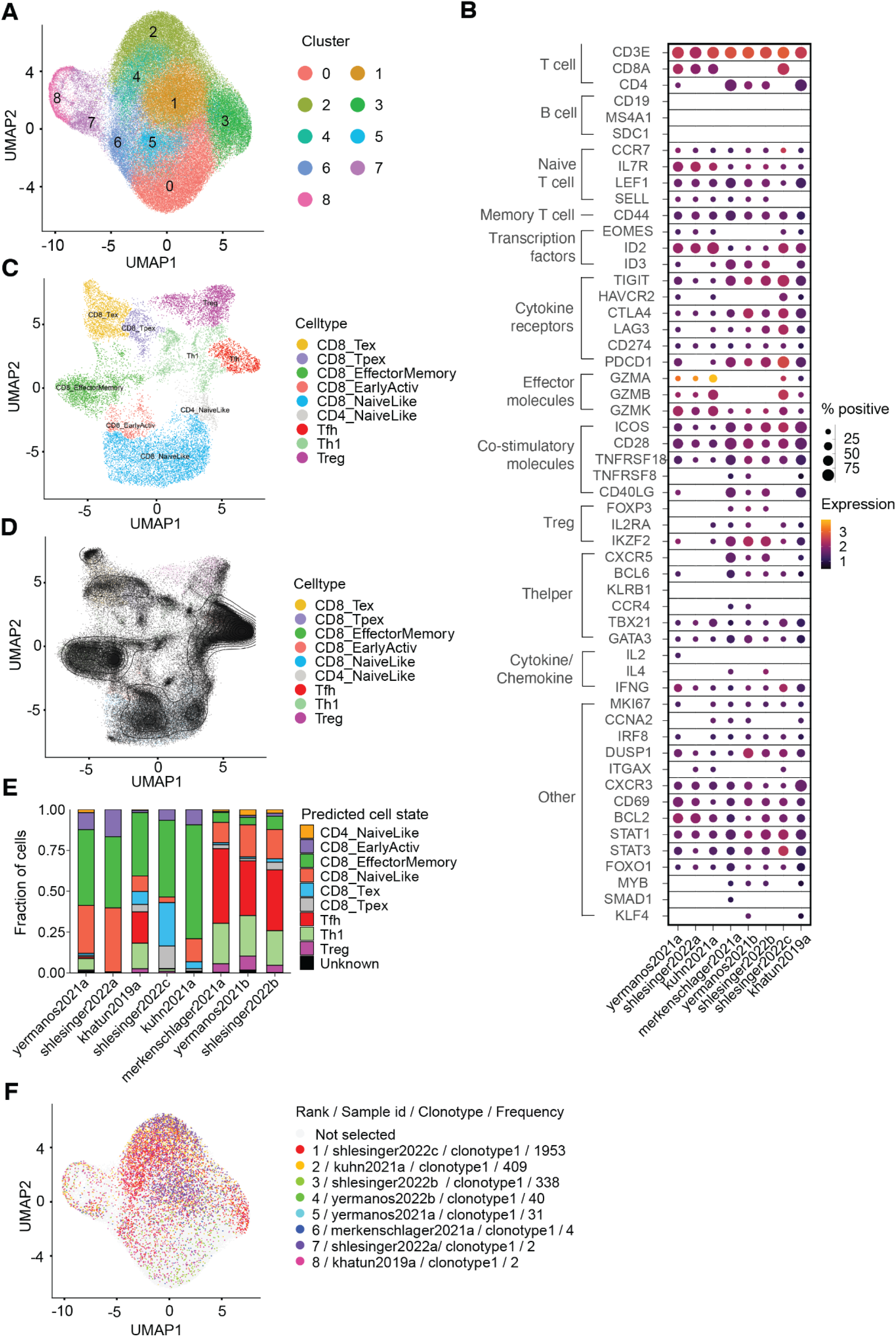
Repertoire and phenotypic breadth of PlatypusDB. A. Uniform manifold approximation projection of murine T cells from various experimental conditions. Each point represents an individual cell and color corresponds to the Seurat assigned cluster. B. Dottile plot showing T cell subset assignment across all datasets based on expression of specific gene lists. C. Projection of reference dataset from projectTILS coloured by T cell phenotype. D. Projection of the complete T cell repertoire (black) over reference dataset from projecTILS (Figure S8C). E. Distribution of T-cell types across experiments from projectTILS. F. Uniform manifold approximation projection of murine T cells from various experimental conditions where each point represents an individual cell. Cells belonging to the most expanded clones overall are highlighted. Clones were defined as containing identical CDRb3+CDRa3 sequences.

**Figure S12.**
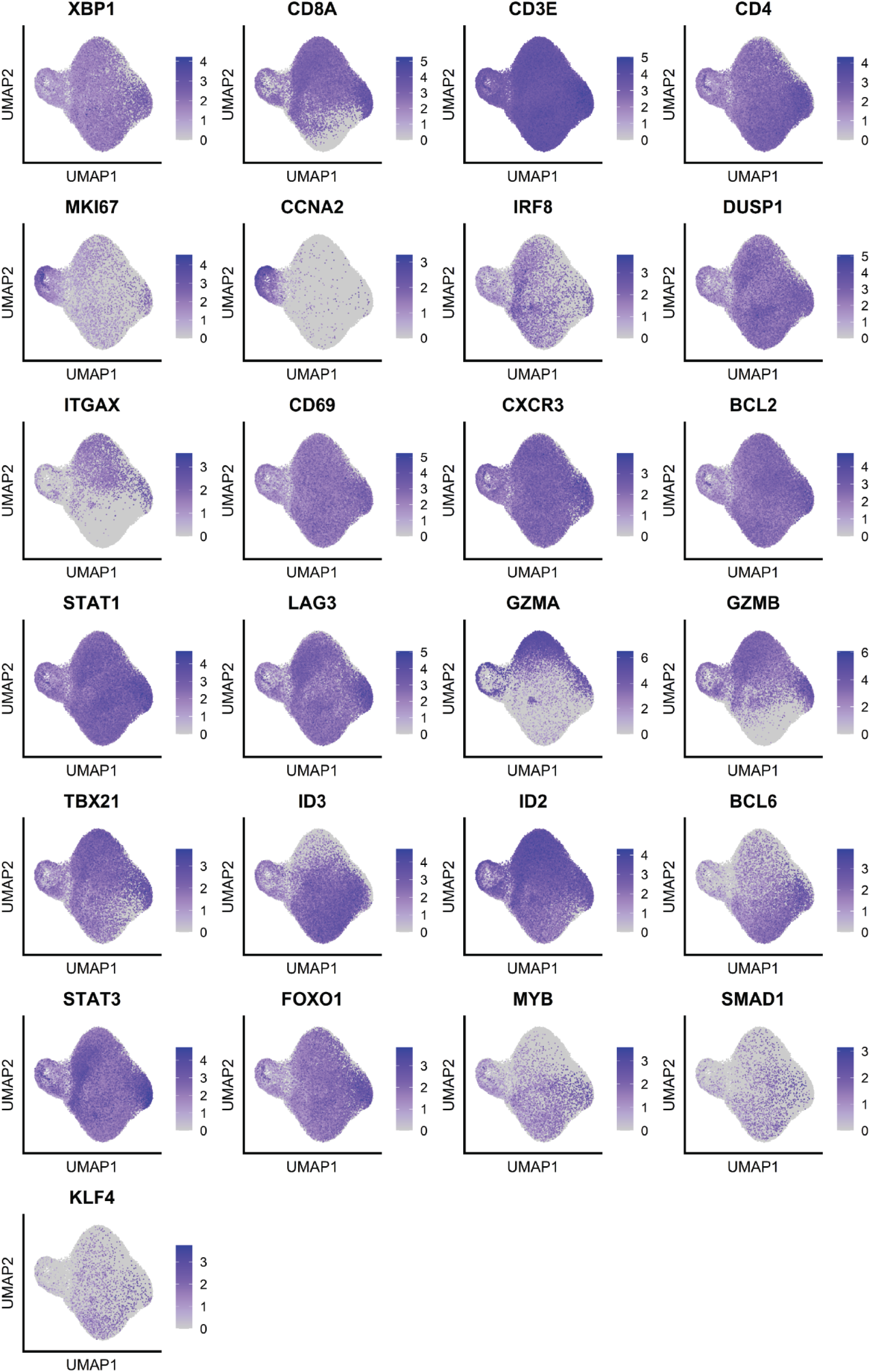
Uniform manifold approximation projection (UMAP) plots showing gene expression for selected T cell associated genes.

**Figure S13.**
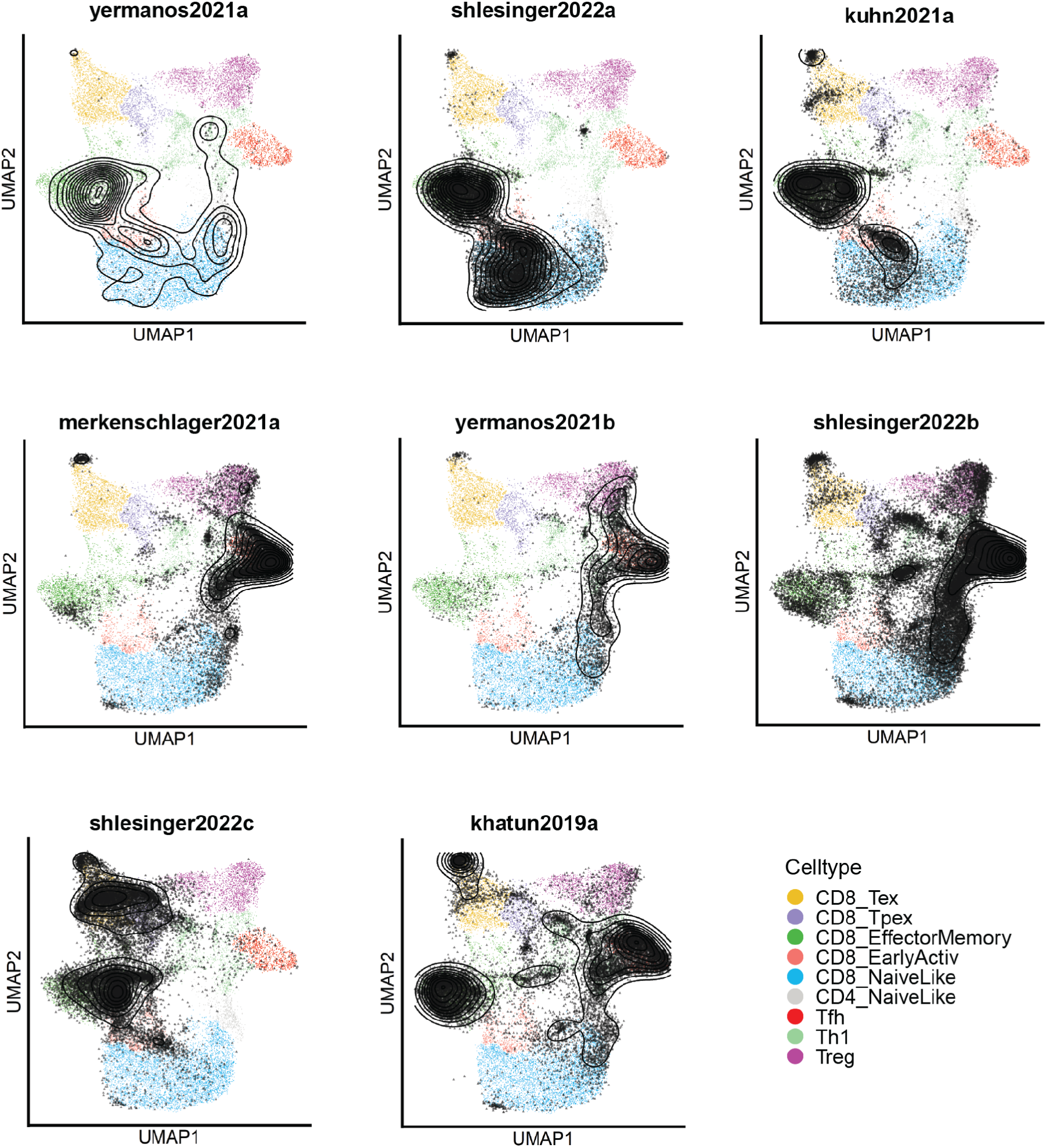
Uniform manifold approximation projection of each individual experiment over the projecTILS reference dataset.

**Figure S14.**
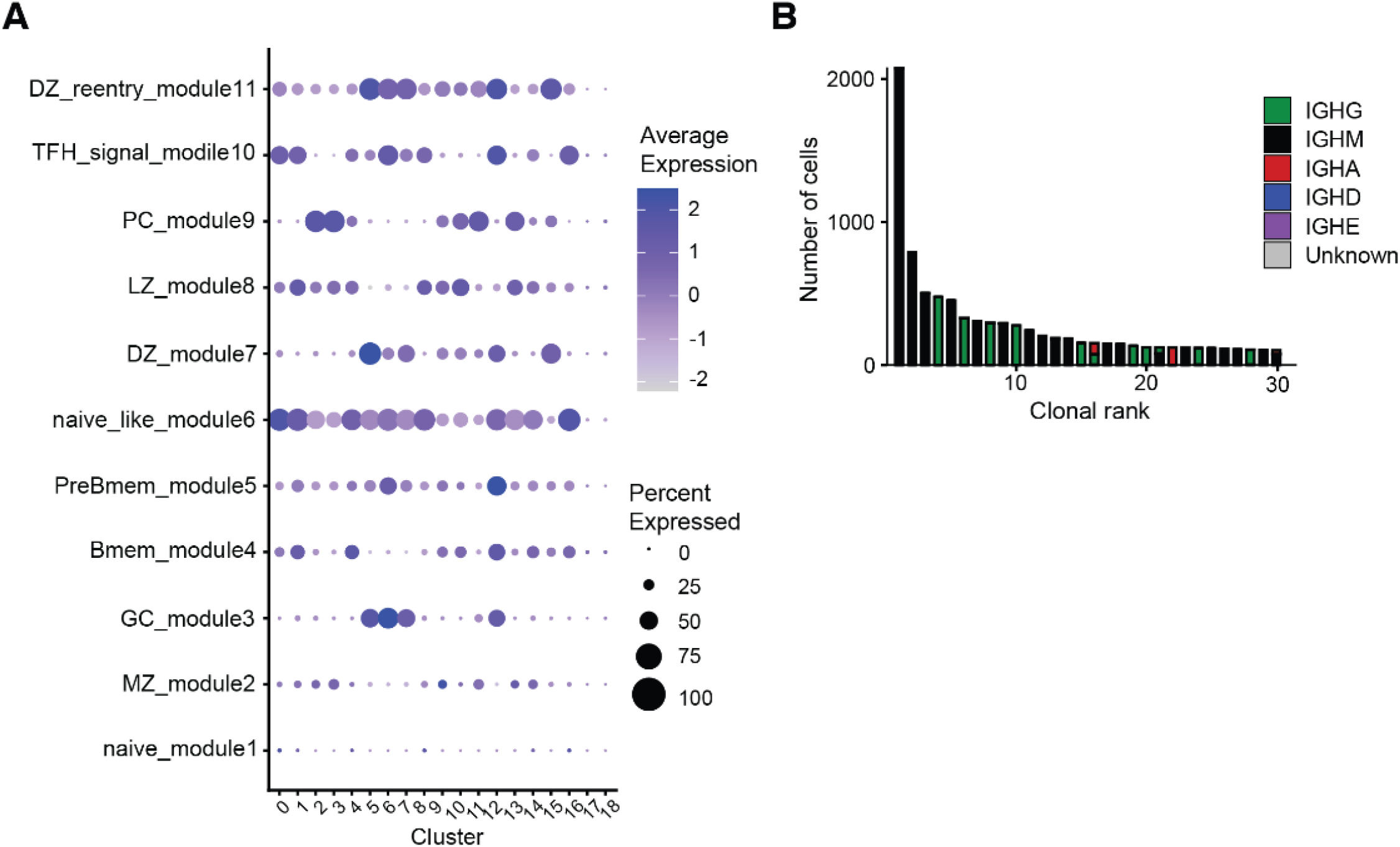
PlatypusDB contains a diverse collection of B cell phenotypes. A. Dottile plot showing B cell subset assignment across all clusters based on expression of specific gene lists (Mathew et al. 2021). B. Analysis of the top 30 most highly expanded clones across all experimental conditions. Clonotyping was performed based on those B cells containing identical HCDR3 + LCDR3 amino acid sequences. The clones were colored by isotype expression.

**Figure S15.**
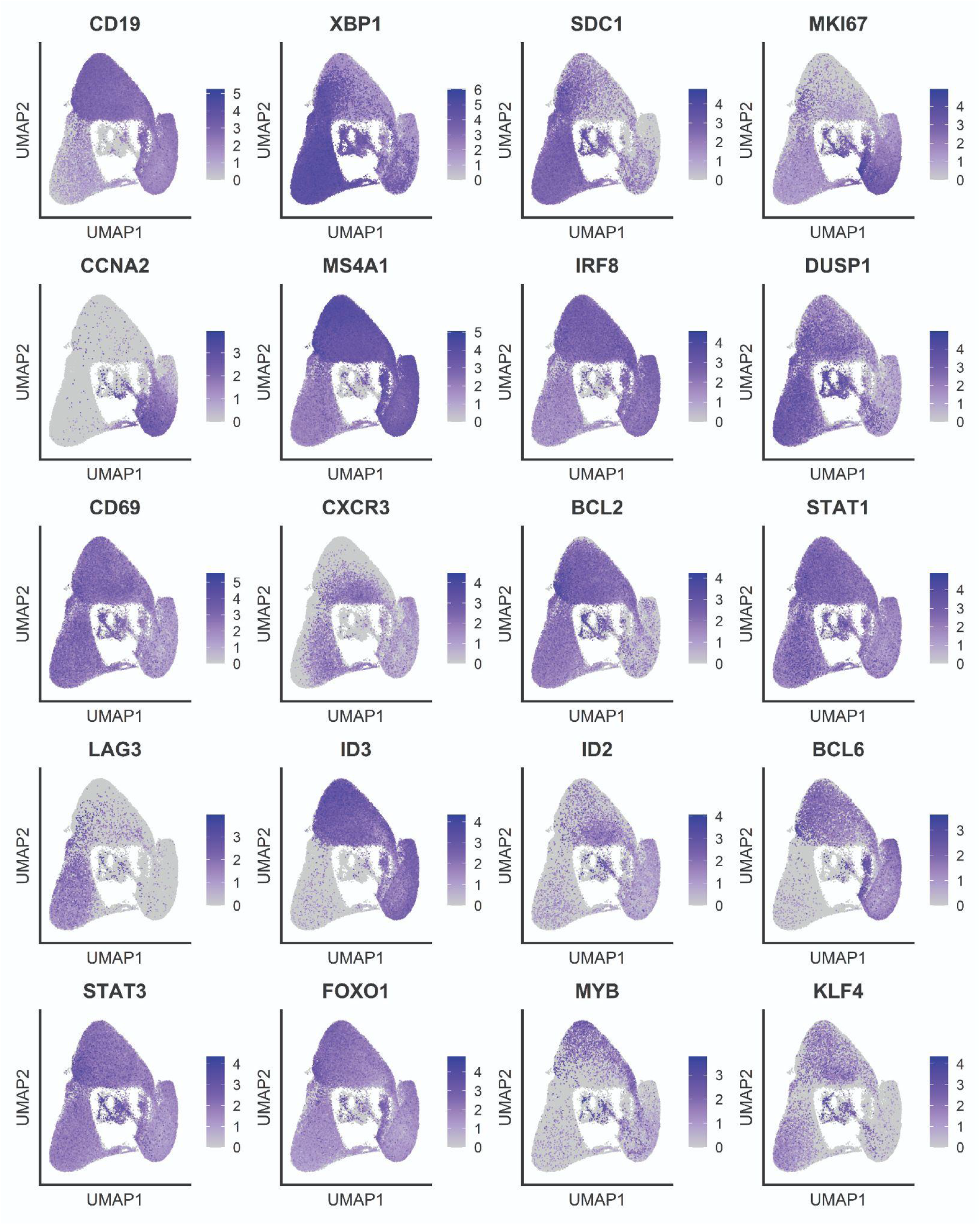
Uniform manifold approximation projection (UMAP) plots showing gene expression for selected B cell genes.

**Table S1.**
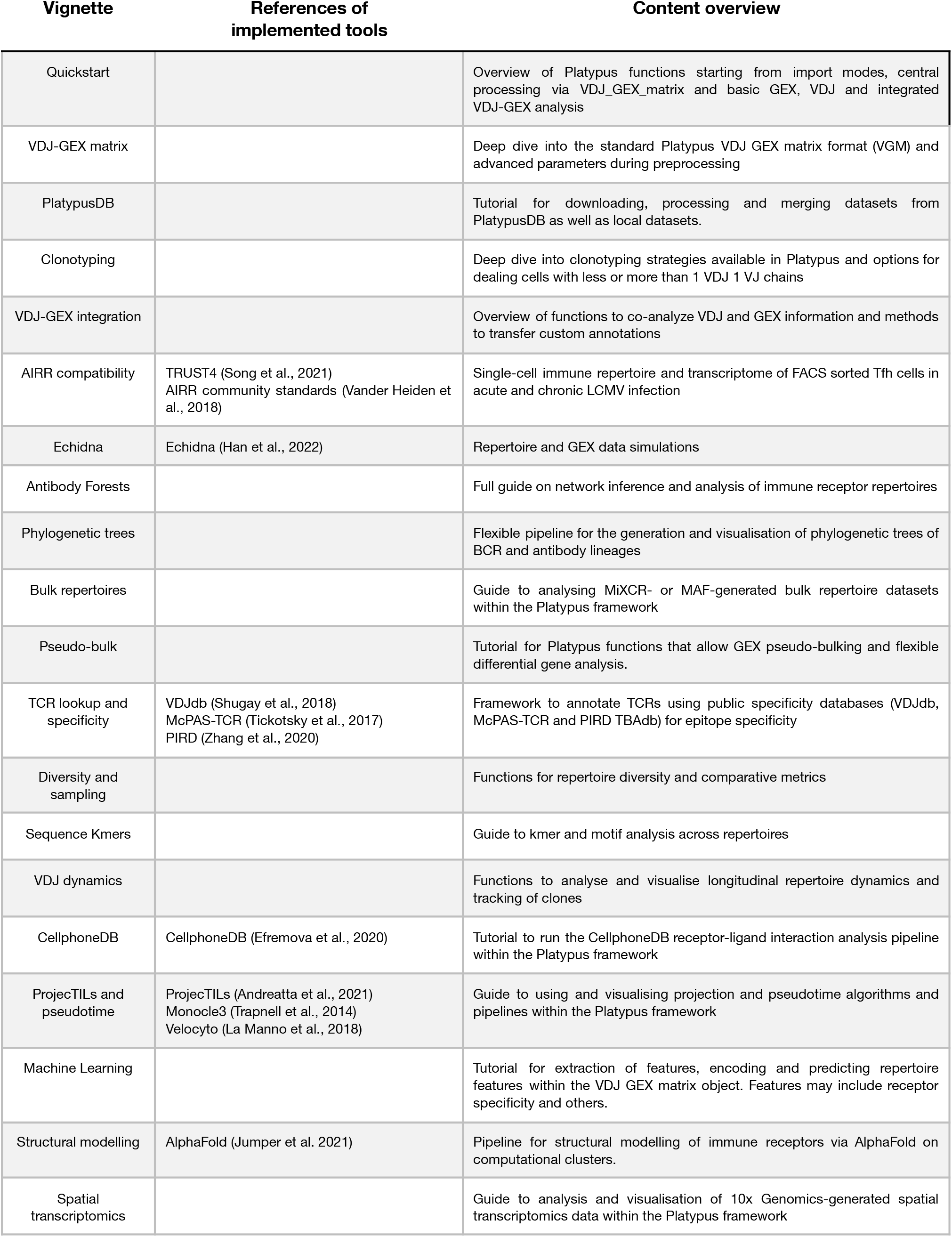
Overview of current tutorials present within the ePlatypus Computational Immunology Ecosystem.

**Table S2.** List of datasets used for the examples and analysis in this manuscript. The PlatypusDB ID is used in code to download datasets. In some cases, more than one dataset was published within the same publication. In that case the title corresponds to the dataset, while the citation and doi correspond to the overarching publication.

| PlatypusDB ID | Citation | DOI | Dataset or publication title |
| --- | --- | --- | --- |
| khatun2019a | Khatun et al., 2021 | 10.1084/jem.20200650 | Single-cell lineage mapping of a diverse virus-specific naive CD4 T cell repertoire |
| kuhn2021a | Kuhn et al., 2022 | 10.3389/fimmu.2022.782441 | Clonally expanded virus specific CD8 T cells acquire diverse transcriptional phenotypes during acute, chronic and latent infections |
| Merkenschlager 2021a | Merkenschlager et al., 2021 | 10.1038/s41586-021-03187-x | Dynamic regulation of Tfh selection during germinal centre reaction |
| mathew2021a | Mathew et al. 2021 | 10.1016/j.celrep.2021.109286 | Temporal dynamics of persistent germinal centers and memory B cell differentiation following respiratory virus infection |
| shlesinger2022a | Shlesinger et al., 2022 | 10.1101/2022.02.07.479381 | Single-cell immune repertoire and transcriptome of GP33+ Tetramer sorted CD8 T cells from single mouse infected with acute LCMV or chronic MCMV infection |
| shlesinger2022b | Shlesinger et al., 2022 | 10.1101/2022.02.07.479381 | Single-cell immune repertoire and transcriptome of FACS sorted Tfh cells in acute and chronic LCMV infection |
| shlesinger2022c | Shlesinger et al., 2022 | 10.1101/2022.02.07.479381 | Single-cell immune repertoire and transcriptome of NP396-tetramer sorted CD8 T cells in a rechallenge déjà vu model |
| yermanos2021a | Yermanos et al., 2021 | 10.1098/rspb.2020.2793 | Single-cell immune repertoire and transcriptome sequencing reveals that clonally expanded and transcriptionally distinct lymphocytes populate the aged central nervous system in mice |
| yermanos2022b | NA | NA | Single-cell immune repertoire and transcriptome sequencing of Tfh cells in Influenza infection |
| kreiner2021a | Shlesinger et al., 2022 | 10.1101/2022.02.07.479381 | Characterization of immune repertoires and phenotypes of T cells in experimental autoimmune encephalomyelitis by single cell sequencing |
| kreiner2021b | Shlesinger et al., 2022 | 10.1101/2022.02.07.479381 | Characterization of immune repertoires and phenotypes of B cells in experimental autoimmune encephalomyelitis by single cell sequencing |
| yewdell2021a | Yewdell et al., 2021 | 10.1016/j.celrep.2021.109961 | Temporal dynamics of persistent germinal centers and memory B cell differentiation following respiratory virus infection |
| neumeier2021a | Neumeier et al., 2021 | 10.1002/eji.202149331 | Single-cell sequencing reveals clonally expanded plasma cells during chronic viral infection produce virus-specific and cross-reactive antibodies |
| neumeier2021b | Neumeier et al., 2021 | 10.1002/eji.202149331 | Single-cell sequencing reveals clonally expanded plasma cells during chronic viral infection produce virus-specific and cross-reactive antibodies |
| agrafiotis2021a | Agrafiotis et al., 2021 | 10.1101/2021.11.09.467876 | B cell clonal expansion is correlated with antigen-specificity in young but not old mice |

## Online Methods

### PlatypusDB architecture

The computational infrastructure for PlatypusDB was developed based on the analysis package Platypus and the Google Cloud storage API. The process of uploading a dataset includes the following steps: First, raw sequencing files are sourced locally, downloaded from public repositories such as GEO, or acquired directly from another research group. Raw reads were then aligned with Cellranger 6.0.1 to the following 10x Genomics reference genomes: refdata-gex-mm10-2020-A, refdata-cellranger-vdj-GRCm38-alts-ensembl-5.0.0, refdata-gex-GRCh38-2020-A, and refdata-cellranger-vdj-GRCh38-alts-ensembl-5.0.0. Raw output files were then uploaded as compressed directories to the PlatypusDB Google Cloud storage database. Raw outputs were then loaded and processed in R resulting in two formats. Firstly a per-sample list object containing main Cellranger output tables and secondly a VDJ-GEX-matrix object from Platypus v3.2.2. This object was generated using the VDJ_GEX_matrix function with default settings, if not otherwise noted. All output objects were uploaded to PlatypusDB using the package googleCloudStorageR. To allow for easy access to the database, download of R objects as well as compressed directories is available directly via URL without the need to install Google Cloud storage compatibility packages for R. The URL for a database lookup table is delivered with Platypus and allows for a single access point to the database, which remains constant as more datasets will be added in the future.

### Data analysis

The filtered feature matrix directory was supplied as input to the VDJ_GEX_matrix function in the R package Platypus (v3.3) (Yermanos, Agrafiotis, et al. 2021), which uses the transcriptome analysis workflow of the R package Seurat (Satija et al. 2015). Only those cells containing less than 20% of mitochondrial reads were retained in the analysis. Genes involved in the adaptive immune receptor (e.g., TRB, TRBV1-1), were removed from the count matrix to prevent clonal relationships from influencing transcriptional phenotypes. Gene expression was normalized using the “harmony” argument in the VDJ_GEX_matrix function. 2000 variable features were selected using the “vst” selection method and used as input to principal component analysis (PCA) using the first 10 dimensions. Graph-based clustering using the Louvain modularity optimization and hierarchical clustering was performed using the functions FindNeighbors and FindClusters in Seurat using the first ten dimensions and a cluster resolution of 0.5. UMAP was similarly inferred using the first ten dimensions. The FindMarkers function from Seurat was used when calculating differentially expressed genes (both across groups or across clusters) with logfc.threshold set to 0 and minimum number of cells expressing each gene set to 0.25 and subsequently supplied to the GEX_volcano function from Platypus. Mitochondrial and ribosomal genes were removed when visualizing DE genes. Feature plots were produced by supplying genes of interest to the function FeaturePlot in Seurat. Module scores for public gene sets (Mathew et al. 2021) were calculated using the AddModuleScore from Seurat. Cells containing no or more than one α/heavy and β/light chain were filtered out for TCR/BCR repertoire analysis. Clones were defined by identical CDR3α/CDRH3 and CDR3β/CDRL3 sequence (nucleotide or amino acid sequence) across all repertoires. Clones represented by more than one cell were considered highly-expanded clones, while single-celled clones were defined as lowly-expanded. The projection of cells onto reference UMAPs and cell state predictions were done using the R package ProjecTILs (Andreatta et al. 2021) under default conditions. Experiments were either individually or all together projected onto the ProjecTILs atlas. For Figures S1 to S8, single-cell immune repertoire sequencing experiments present in PlatypusDB were formatted into a single VDJ_GEX_matrix object that was then supplied to downstream analyses pipelines. Specifically, the pseudobulk analysis was performed using the pseudo_bulk_DE function from Platypus. Sampling and diversity analyses were performed using the VDJ_diversity and VDJ_rarefaction. The sequence similarity network of clusters was generated using the AntibodyForests_communities function and then colored using the AntibodyForests_plot. Node and edge metrics were calculated using the AntibodyForests_metrics function from Platypus. The PlatypusML_classification function from Platypus, which takes input the encoded features obtained from the PlatypusML_extract_features function, was used to run cross validation on a specified number of folds for different classification models (XGBoost, SVM, Random Forest, Logistic Regression & Gaussian Naive Bayes), outputting the AUC scores, ROC curve and confusion matrix for each classification model. Prediction of receptor-ligand interaction was calculated on the single-cell gene expression data using the CellPhoneDB software. The heatmap and dot plot were generated as an output of the CellPhoneDB_analyse function from Platypus. Receptor structures in the Steropodon workflow were obtained using the Steropodon_model function and visualized using the Steropodon_visualize function in Platypus.

### Data visualization

Figure 1 and the supplementary graphical overviews were created with Biorender.com. Feature plots were produced using “FeaturePlot” (Seurat 4.0). Volcano plots were produced using “GEX_volcano” (Platypus v3.3). Dottile plots were produced using DotPlot (Seurat 4.0). All other figures were produced using Prism v9 (Graphpad).

## Data availability

The accession numbers and publications for the sequencing data used in this manuscript are located in table S1. Platypus code used in this manuscript can be found at github.com/alexyermanos/Platypus.

## Acknowledgements

We acknowledge and thank Dr. Christian Beisel, Elodie Burcklen, Ina Nissen, and Mirjam Feldkamp at the ETH Zurich D-BSSE Genomics Facility Basel for excellent support and assistance. We also thank Nathalie Oetiker and Franziska Wagen for excellent experimental support.

## Funding

This work was supported by the ETH Zurich Research Grants (to STR and AO), an ETH Seed Grant (AY) and support from the “la Caixa” Foundation (ID 100010434, fellowship code: LCF/BQ/EU20/11810041) to MMC.

## Competing Interests

There are no competing interests.

## Notes

https://alexyermanos.github.io/Platypus/index.html

### Competing Interest Statement

The authors have declared no competing interest.

